# Opponent visuospatial coding structures responses during memory recall and visual perception in medial parietal cortex

**DOI:** 10.1101/2024.09.13.612875

**Authors:** Catriona L. Scrivener, Edward H. Silson

## Abstract

The mechanisms linking perceptual and memory representations in the brain are not yet fully understood. In early visual cortex, perception and memory are known to share similar neural representations, but how they interact beyond early visual cortex is less clear. Recent work identified that scene-perception and scene-memory areas on the lateral and ventral surfaces of the brain are linked via a shared but opponent visuospatial coding scheme, suggesting that shared visuospatial coding might provide a framework for perceptual-memory interactions. Here, we test whether the pattern in visuospatial coding within category-selective memory areas of medial parietal cortex structures responses during memory recall and visual perception. Using functional magnetic resonance imaging, we observe signatures of visuospatial coding in the form of population receptive fields (pRFs) with both positive and negative response profiles within medial parietal cortex. Crucially, the more dissimilar the timeseries of a pair of positive/negative pRFs within a region, the more dissimilar their responses during both memory recall and visual perception - tasks that place very different demands on these regions: internally oriented memory recall versus externally oriented visual perception. These data extend recent work to suggest that the interplay between pRFs with opponent visuospatial coding may play a vital role in integrating information across different representational spaces.

## Introduction

Interaction with the complex visual world around us requires the constant integration of perceived visual information with the retrieval of previously encoded information from memory. Understanding how perceptual and memory representations interact is therefore an enduring puzzle in both psychology and neuroscience (Dijkstra et al., 2020; Favila et al., 2020; Groen et al., 2021; Steel et al., 2024). Prior computational (Marr, 1971; Norman & O’Reilly, 2003), and neuroimaging work (Klein et al., 2004; Slotnick, 2009; Thirion et al., 2006; Polyn et al., 2005) suggests that perception and memory may share similar neural representations, as the recall of previously encoded stimuli recruits or reactivates retinotopically specific portions of early visual cortex (i.e., neural reinstatement: Klein et al., 2004; Slotnick, 2009; Thirion et al., 2006).

However, how perception and memory representations integrate beyond early visual cortex is less clear. For example, although visual cortex is known to encode perceptual information within a visuospatial reference frame, according to locations in the visual field relative to one’s gaze position (Groen et al., 2021; Wandell et al., 2007), anterior memory-driven regions (e.g., hippocampus) are usually considered to be governed by more abstract coding schemes such that visuospatial information is abstracted away as information propagates through the cortical hierarchy (Guclu & van Gerven, 2015; Margulies et al., 2016; Popham et al., 2021a). Thus, how perception and memory representations are integrated beyond early visual cortex is a matter of current debate.

Work spanning the last decade demonstrates that the visuospatial coding scheme is more extensively represented in the brain than previously thought and may offer a plausible framework for such perceptual-memory integration. For instance, there are over twenty-five separate visual field maps identified throughout the dorsal and ventral visual streams (Amano et al., 2009; Arcaro et al., 2009; Brewer et al., 2005; Larsson & Heeger, 2006; Wandell et al., 2007; Wang et al., 2015), and signatures of visuospatial coding, such as biases for the contralateral visual field, and spatially-sensitive population receptive fields (pRFs, Dumoulin & Wandell, 2008) have been identified within category-selective regions of visual cortex - once thought position invariant (Silson et al., 2015; Silson, Groen, et al., 2016; Silson, Steel, et al., 2016; Silson, Groen, et al., 2021), the default-mode network (Szinte & Knapen, 2020) and even hippocampus (Knapen, 2021; Silson, Zeidman, et al., 2021). Together, this body of work suggests that the brain retains the visuospatial coding scheme throughout the cortical hierarchy and may make use of it to aid different cognitive processes (Groen et al., 2021; Szinte & Knapen, 2020).

Very recent work explored this possibility within regions involved in the perceptual processing and mental recall of scenes, finding shared access to the same visuospatial coding scheme (Steel et al., 2024). Specifically, it was shown that whereas pRFs in scene-perception areas (Occipital Place Area [OPA] and Parahippocampal Place Area [PPA], Dilks et al., 2013; Epstein & Kanwisher, 1998) had positive response profiles **(+ve pRFs: increased response to a stimulus within the receptive field)**, a significant percentage of pRFs in their paired scene-memory areas (Lateral Place Memory Area [LPMA] and Ventral Place Memory Area [VPMA], Steel et al., 2021, 2024) had negative response profiles **(−ve pRFs: decreased response to a stimulus within the receptive field)**. Moreover, during memory recall, there was a push-pull relationship between these two sets of pRFs between regions, whereby when activity of +ve pRFs was high in perceptual regions (i.e., OPA/PPA), activity of −ve pRFs was low in memory regions (i.e., LPMA/VPMA) and vice-versa (Steel et al., 2024). This finding suggests that these coupled scene-perception and scene-memory areas make use of the same visuospatial coding scheme, but with opponent profiles, in order to represent bottom-up sensory information and top-down memory information while limiting interference.

Whilst this prior work represents a novel observation, important considerations are that this process was: **A)** Only tested within perception and memory areas on the lateral and ventral surfaces of the brain and **B)** was only explored within the context of scene-perception and scene-memory, and not other stimulus categories like people. Crucially, prior work has identified additional regions of the brain that are critical to memory recall (Silson, Gilmore, et al., 2019; Silson, Steel, et al., 2019). Indeed, within the medial parietal cortex (MPC) there exists an alternating pattern of four regions selectively engaged during the mental recall of either people or places (**Figure 1**). So, while it has been shown that scene-perception and scene-memory areas on the lateral and ventral surfaces are linked via visuospatial coding, it is currently unknown whether such an organisation persists into people and place memory regions in MPC.

**Figure 1:**
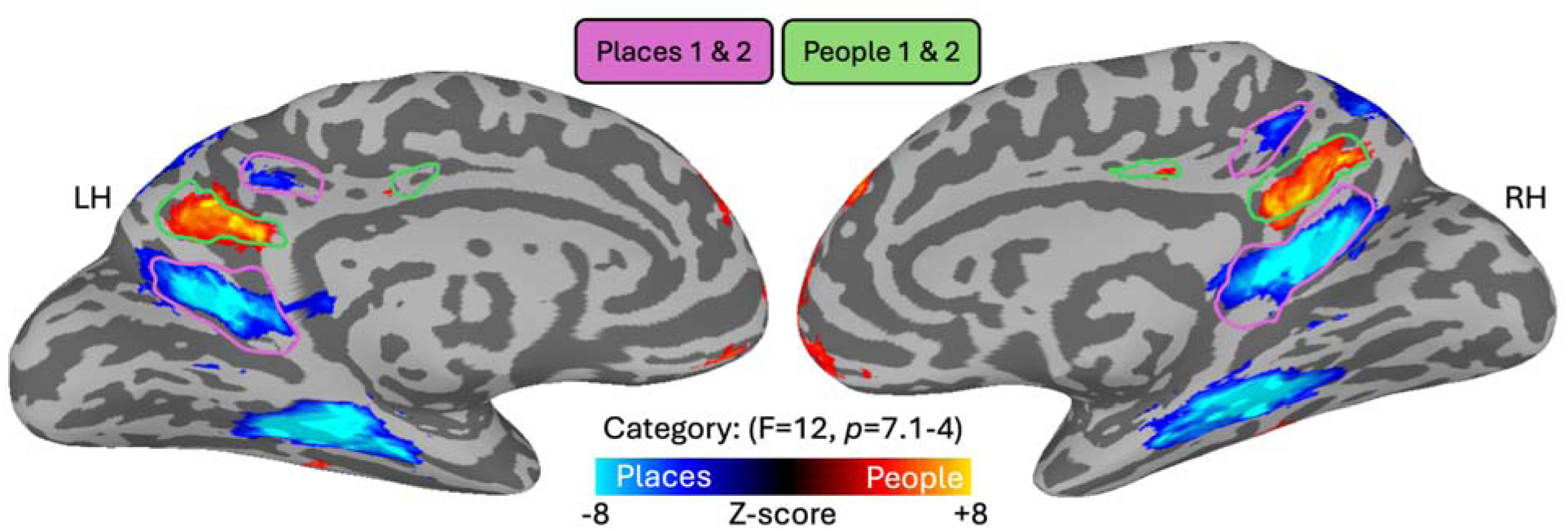
People and place memory regions in medial parietal cortex. Group level results from a whole-brain linear mixed model contrast for the recall of people versus place stimuli. The results are displayed on partially-inflated cortical surfaces of a sample subject, separately for the left and right hemispheres. Activation was thresholded with a minimum F value of 12 for the main effect of category (p < 7.1^−4^). Regions with greater activation for places are displayed in cold colours, and those with greater activation for people displayed in hot colours. Place-selective regions (Places1, Places2) from prior work (Silson, Steel, et al., 2019) outlined in purple, and people-selective regions (People1, People2) outlined in green show a high-degree of overlap with the current ROIs.

Here, we asked whether people and place memory regions of MPC also show signatures of visuospatial coding (i.e., +ve and −ve pRFs). We took advantage of two existing datasets from our lab containing a mixture of pRF mapping (Dataset1 and 2, n=36), scene-face localisers (Dataset1 and 2, n=36), scene perception (Dataset1, n=24), and memory recall data (Dataset2, n=12). We had two main goals: **1)** Quantify the presence of visuospatial coding throughout MPC using pRF modelling, and **2)** Examine the extent to which the pattern of visuospatial coding between +ve and −ve pRFs structures responses across different cognitive processes: memory recall and visual perception. To preview, we show that memory recall regions of MPC exhibit visuospatial coding and contain significant proportions of −ve pRFs. Crucially, we show that the pattern in visuospatial coding between +ve and −ve pRFs structures the responses across different cognitive tasks, such that the more dissimilar the timeseries between of pair of positive/negative pRFs within a region, the more dissimilar their responses during both memory recall and visual perception.

## Methods

### Dataset descriptions

Here we used two independent fMRI datasets collected by our team at the University of Edinburgh Imaging Facility RIE, Edinburgh Royal Infirmary. Both MRI datasets were acquired using the same Siemens 3T Prisma scanner and 32 channel head coil and very similar sequence parameters. Independent results from Dataset1 can be found in Scrivener et al., (2024, in press).

Dataset1 had 24 participants (mean age: 24, 7 males), and Dataset2 had 12 participants (mean age: 27, 3 males). Participants included students from the University of Edinburgh and individuals from the surrounding areas. Participants had normal or corrected-to-normal vision, and were free from neurological or psychiatric conditions. Written consent was obtained from all participants in accordance with the Declaration of Helsinki and a consent form approved by the School of Philosophy, Psychology and Language Sciences ethics committee of the University of Edinburgh. Participants were given monetary compensation for their time.

For both datasets, participants completed 2 hours of scanning with a break in between (ranging from 15-60 minutes, depending on scanner timetabling constraints). In the first hour of scanning for both datasets we acquired T1 and T2 weighted structural scans, followed by functional runs of population receptive field (pRF) mapping (Dumoulin & Wandell, 2008), lasting six minutes each (Dataset1=6 runs per participant, Dataset2=4 runs). In the second hour of scanning for both datasets we first acquired 2 runs of a scene-face localiser scan lasting 5 minutes each. In Dataset1 we next acquired 2 runs of an event-related visual perception task, lasting 8 minutes each. In Dataset2 we instead acquired 6 runs of a recall task, lasting 6 minutes each. One participant only completed 4 of the 6 recall runs.

Using both datasets we have a mixed sample of pRF mapping (both datasets, n=36), scene-face localisers (both datasets, n=36), scene perception (Dataset1, n=24), and memory recall data (Dataset2, n=12).

### MRI recording parameters

For both datasets we acquired two structural images; T1 weighted (TR = 2.5s, TE = 4.37ms, flip angle = 7 deg, FOV = 256mm x 256mm x 192mm, resolution = 1mm isotropic, acceleration factor = 3), and T2 weighted (TR = 3.2s, TE = 408ms, flip angle = 9 deg, FOV = 256mm x 240mm x 192mm, resolution = 0.9mm isotropic, acceleration factor = 2). The majority of the functional scans across both datasets were acquired using the same multiecho multiband echo planar imaging sequence with the following parameters (TR = 2, TEs = 14.6ms, 35.79ms, 56.98ms, MB factor = 2, acceleration factor = 2, 48 interleaved slices, phase encoding anterior to posterior, transversal orientation, slice thickness = 2.7mm, voxel size = 2.7mm x 2.7mm, distance factor = 0%, flip angle = 70 degrees). The first 6 participants in Dataset1 were recorded with a larger number of slices (52) and slightly different echo times (14.6ms, 32.84ms, 51.08ms). To accommodate larger heads, we reduced the number of slices from 52 to 48 from participant 6 onwards, which provided greater coverage in the anterior-posterior direction. Although we were able to achieve whole-brain coverage for participants 1-5 who had smaller head sizes, we made this change as a precaution.

### Stimuli and tasks

#### Population receptive field modelling

During pRF mapping sessions, a bar aperture traversed gradually through the visual field, whilst revealing randomly selected scene fragments from 90 possible scenes. During each 36s sweep, the aperture took 18 evenly spaced steps every 2s (1 TR) to traverse the entire screen. Across the 18 aperture positions all 90 possible scene images were displayed once. A total of eight sweeps were made during each run (four orientations, two directions). Specifically, the bar aperture progressed in the following order for all runs: Left to Right, Bottom Right to Top Left, Top to Bottom, Bottom Left to Top Right, Right to Left, Top Left to Bottom Right, Bottom to Top, and Top Right to Bottom Left. The bar stimuli covered a circular aperture (diameter = 12 degrees of visual angle). Participants performed a colour detection task at fixation, indicating via button press when the white fixation dot changed to red. Colour fixation changes occurred semi-randomly, with approximately two-colour changes per sweep (Silson et al., 2015).

#### Scene-face localiser

During each run, colour images of scenes and faces were presented at fixation (10×10 degrees of visual angle) in 16s blocks (20 images per block; 300ms per image, 500ms blank). Participants responded via an MRI compatible button box whenever the same image appeared sequentially (randomly occurring twice per run).

#### Scene perception: Dataset1

During two event-related scene perception scans, participants were presented with 96 complex scenes (12×9 degrees of visual angle, 500ms each) in a randomised order as in (Kravitz et al., 2011). Interstimulus intervals (3-7s) were chosen to optimise the ability of the later deconvolution to extract the responses to each scene. Participants fixated centrally and performed an orthogonal fixation colour-detection task, pressing a button (via MRI compatible button box) every time the green fixation cross turned red (randomly occurring 9 times per run).

#### Recall: Dataset2

Before attending their first session, participants in Dataset2 were asked to provide the researchers with images of six personally familiar people and six personally familiar places that they were confident they could recall easily. We asked participants to save these images with a name that would serve as a cue for them to recall the stimuli, such as ‘Sophie’, or ‘Office’. During each trial, participants were presented with a randomly chosen word cue for 500ms, followed by 9.5s of a green fixation cross during which they were asked to create a mental image of the cued stimulus. Each recall period was followed by a jittered ITI of 2.5-7s where a white fixation cross was presented. Each recall cue was presented twice per run across 6 runs (12 total repetitions). We specifically asked participants to generate images that closely matched the pictures that they had provided to make the recall as similar as possible across repetitions.

#### Pre-processing

MRI scans were processed using AFNI (Cox, 1996), Freesurfer, and SUMA (Saad & Reynolds, 2012). Dummy scans were removed from the start of each run (3dTcat). Slice time correction was then performed (3dTshift), aligning each slice with a time offset of zero. The skull was removed from the first echo 1 scan and used to create a brain mask (3dSkullStrip and 3dAutomask). The first echo 2 scan was used as a base for motion correction and registration with the T1 structural scan (3dbucket). Motion parameters were estimated for the echo 2 scans (3dVolreg) and applied to the other echos (3dAllineate). After completing the standard pre-processing, the data was also processed using tedana (Evans et al., 2015; Kundu et al., 2012) to denoise the multi-echo scans (version 0.0.12, using default options). The tedana optimally combined and denoised output was then scaled so that each voxel had a mean value of 100 (3dTstat and 3dcalc). This means that the fMRI values can be interpreted as a percentage of the mean signal, and effect estimates can be viewed as percentage change from baseline (Chen et al., 2017). For the pRF data, an average of the runs was then taken to leave a single timeseries for further analysis.

The session 1 structural scans were aligned to the functional data collected in session 1 (align_epi_anat) with a multi-cost function (including LPC, MNI, and LPA) and manually checked for accuracy. For almost all subjects we used the default LPC method output (localised Pearson correlation), as this worked sufficiently well. For three subjects in Dataset1 we used the NMI output (normalised mutual information). Functional data collected in session 2 was aligned to the session 1 structural to ensure that all functional data had the same alignment. Freesurfer reconstructions were estimated using both the T1 and T2 scans (recon-all) from session 1, and the output used to create surfaces readable in SUMA (SUMA_Make_Spec_FS). The SUMA structural was then aligned to the session 1 experimental structural to ensure alignment with the functional images (SUMA_AlignToExperiment). Surface based analyses were conducted using the SUMA standard cortical surface (std.141).

#### pRF modelling and amplitude definition

Population receptive fields were estimated using AFNI’s non-linear fitting algorithm (3dNLfim) and the GAM basis function. Full details are provided elsewhere (Silson et al., 2015). Suprathreshold pRFs were defined as those with R² > 0.08, as in prior work (Steel et al., 2024). pRFs with an estimated size larger than the visual presentation were also excluded (sigma > .95). Visuospatial organisation refers to regions with spatially specific visual responses in the absence of traditionally defined retinotopic maps (e.g. polar angle and eccentricity maps in early visual cortex). These regions may have many suprathreshold pRFs, clearly indicating that their response varies with the spatial position of the stimulus, but do not have visible gradients along which to draw retinotopic maps (Groen et al., 2021). pRFs with an amplitude greater than 0 were defined as positive (+ve), whereas pRFs with an amplitude less than 0 were defined as negative (−ve). We calculated the percentage of suprathreshold −ve pRFs within each ROI and entered these values (averaged across hemispheres) into a linear mixed model (LMM) with Category (People, Places) and Position (Posterior, Anterior) as factors. LMM’s were conducted in R (v1.3; lme4 v27.1; lmerTest v3.1). We also computed the visual field coverage of each ROI, which indicates the locations within a visual field that evoke the greatest response across voxels. We calculated coverage as the best Gaussian population receptive field model for each suprathreshold node within an ROI. We used a max operator that reflects the maximum value from all pRFs within the ROI for each point in the visual field (Winawer et al., 2010).

### Recall univariate activation

The activity associated with each stimulus in the recall scans was deconvolved using a BLOCK basis function (BLOCK(9.5,1), 3dDeconvolve and 3dREMLfit) aligned to the onset of the recall period. Each run was modelled separately and estimates of subject motion were included as regressors of no interest. The output of the model was projected onto the SUMA standard cortical surface (3dVol2Surf). Group level analysis was conducted on the surface data using a linear mixed model (3dLME). We included subject as a random effect and modelled the main effect of category (people vs. places), the main effect of run (runs 1 to 6), and an interaction between category and run effects.

### Defining medial parietal regions of interest

Medial parietal ROIs were taken from previous work (Silson, Steel, et al., 2019) to avoid issues of circularity in our analysis (Kriegeskorte et al., 2009). Here, two ROIs on the medial parietal surface with increased BOLD activation during recall of place stimuli were named as Places1 and Places2. A further two ROIs with greater activation for people stimuli were named People1 and People2. To compare the spatial location of prior ROIs with the group-level recall data from Dataset2, we calculated the dice-coefficient of overlap between both sets of ROIs, which is given as: (2*Overlap between ROIs / size of ROI A + size ROI B). The independent ROIs from prior work were transformed from surface space into each individual participants volumetric space using the same parameters that were used to project their data to the surface (3dSurf2Vol). All subsequent analyses were conducted using the voxels within these volumetric ROIs.

### Visual perception univariate analysis

The activity associated with each stimulus in the visual perception scans was deconvolved using a GAM basis function aligned to the onset of each stimulus (3dDeconvolve and 3dREMLfit). The two runs were modelled together and each stimulus regressor included two onsets.

### Representational dissimilarity analysis

Representational dissimilarity analysis was conducted in the volume within each MPC ROI in the left and right hemispheres separately (Places1, Places2, People1, and People2). In the first analysis (n=36), we tested the hypothesis that there was an inverse relationship between the visual field coverage of each pRF (estimated using the x, y, and size parameters from pRF modelling) and the average timeseries during pRF mapping. That is, the more similar the coverage across a pair of +ve and −ve pRFs within an ROI, the more dissimilar their timeseries during pRF mapping. This is based on the principle that activity within a +ve pRF increases when stimuli are presented in its preferred spatial location, whereas activity within a −ve pRF decreases. To test this hypothesis, for each participant and ROI we constructed a representational dissimilarity matrix (RDM) of the pairwise dissimilarity (1-Pearson’s r) between the visual field coverage for all pairs of +ve and −ve pRFs (pdist2, MATLAB). We then correlated this coverage RDM with a similar RDM constructed using the pRF timeseries (again across +ve and −ve pRFs). To test whether there was a significant negative correlation between these RDMs across participants, we entered the normalised correlation values (Fisher’s transform, atanh, MATLAB) into a one-sample t-test (two-tailed).

Next, we asked whether the similarity in visuospatial coding between +ve and −ve pRFs (i.e., pRF timeseries RDM) was also present during memory recall (n=12). To do this we correlated the pRF timeseries RDM (as described above) with a similar RDM constructed using the recall timeseries, hypothesising that there would be a positive correlation. For each participant we computed one RDM per recall run before using the average RDM in the correlation analysis. We tested for a significant positive correlation across participants using the normalised correlation values, as above.

Lastly, we tested whether the visuospatial coding present in our ROIs also structured the responses during visual perception (n=24). For this, we correlated the pRF timeseries RDMs with RDMs constructed using the scene-perception timeseries. We again tested for significant positive correlations across participants using the normalised correlation values, as above.

#### Control analyses

We considered whether our results could be explained by other less interesting factors such as differences in temporal signal-to-noise ratio (tSNR). tSNR was calculated as the mean timeseries in each voxel divided by its standard deviation across time. As this calculation gives one value per voxel, we constructed tSNR RDMs using the Euclidean distance between +ve and −ve pRFs for both the pRF and recall timeseries. If the relationship observed between the pRF RDMs and recall RDMs could be explained by tSNR in the data, we expected to find positive correlations between the tSNR RDMs across pRF and recall data.

As a final control test, we shuffled the +ve/−ve labels of the voxels before computing the correlation between pRF and recall RDMs. Shuffling the pRF labels and therefore the timeseries removes the existing relationships between the timeseries and destroys the similarity in visuospatial coding. We therefore expected that unless other extraneous factors were driving their relationship, shuffling would result in a non-significant correlation between the two RDMs.

## Results

Here, we focus on quantifying the presence of visuospatial coding within people and place memory regions of MPC. Prior work identified an alternating pattern of people and place memory recall regions within MPC along the posterior-anterior axis (Places1, People1, Places2 & People2, **see Figure 1**). As such, we initially sought to replicate this finding by comparing the magnitude of activity during people versus place recall (n=12, **see Methods**). At the group-level, we were able to identify all four regions-of-interest (ROIs) in both hemispheres. Moreover, the spatial location and topography of these ROIs was entirely consistent with prior work (Dice-coefficient of overlap between prior and current ROIs; LH: Places1=0.86, People1=0.56, Places2=0.89, People2=0.09; RH: Places1=0.86, People1=0.66, Places2=0.84, People2=0.46). Given the high-degree of similarity between the two datasets and to maximise the power in our sample and avoid issues of circular analyses (Kriegeskorte et al., 2009), all subsequent analyses made use of ROIs from independent prior work (Silson, Steel, et al., 2019).

### Visuospatial coding in people and place memory areas of MPC

Next, we examined whether people and place memory areas of MPC showed signs of visuospatial coding using pRF modelling. Here, we combined data from two separate pRF datasets acquired with the same experimental paradigm (Dataset1 n=24, Dataset2 n=12, **see Methods**). For consistency with prior work, we calculated the percentage of suprathreshold (R^2^>0.08) −ve pRFs within each ROI (**Figure 2**). All people and place memory regions of MPC contained significant percentages of −ve pRFs (*t*-test versus zero, all *t*-values >15.13, all *p*-values <0.001, **see Table S1 for full statistical breakdown**), which did not differ between hemispheres (*t*(245)=1.95, *p*=0.05). The prevalence of −ve pRFs in MPC, appeared relatively constant along the posterior-anterior axis. This consistency is in contrast with prior work in which the prevalence of −ve pRFs was shown to increase from scene-perception to scene-memory areas on both the lateral (i.e., OPA to LPMA) and ventral (i.e., PPA to VMPA) surfaces (Steel et al., 2024). Further, the prevalence of −ve pRFs in MPC were significantly higher than those in LPMA/VMPA on average (*t*(35)=6.88, *p*=5.36-8).

**Figure 2:**
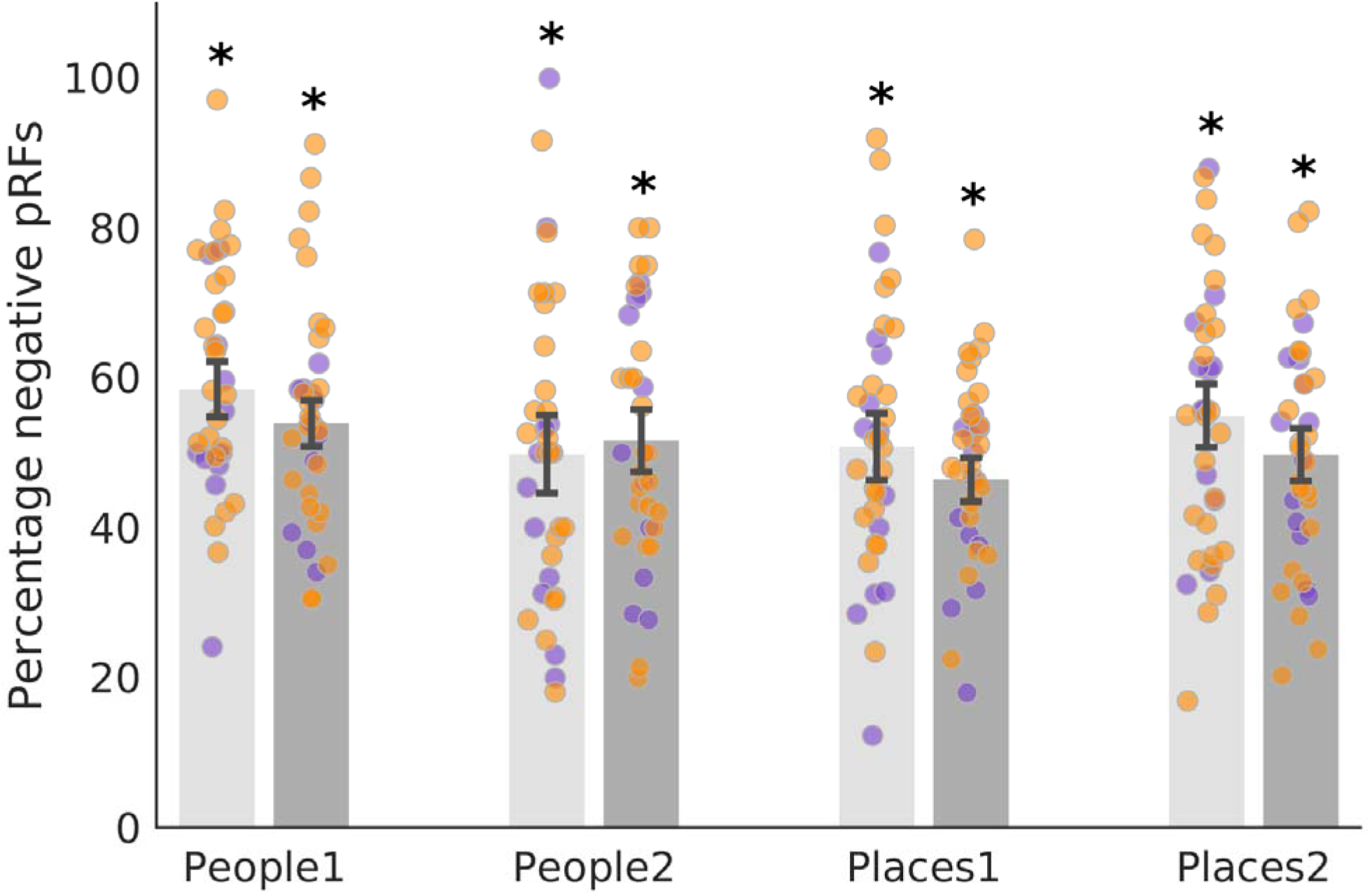
Visuospatial coding within people and place regions of MPC. Bars represent the mean percentage of suprathreshold (R²>0.08) −ve pRFs in each region (left hemisphere=light grey, right hemisphere=dark grey). Individual participant data points are included from both datasets (Dataset1 = orange dots, Dataset2 = purple dots). Error bars represent the standard error of the mean (SEM). Each ROI contains a significant percentage of −ve pRFs (\**p*<0.001).

To quantify these effects, we submitted these values (averaged across hemispheres) into a linear mixed model (LMM) with Category (People, Places) and Position (Posterior, Anterior) as factors. Neither the main effect of Position (F(1, 245)=1.80, *p*=0.18), nor Category (F(1, 245)=3.68, *p*=0.06) were significant. These effects were qualified, however, by a significant Category by Position interaction (F(1, 245)=6.70, *p*=0.01), which reflects higher prevalence of −ve pRFs posteriorly for people (i.e., People1 > People2), but anteriorly for places (i.e., Places2 > Places1), respectively. People and place memory areas within MPC had significant levels of −ve pRFs, but importantly they were not universally negative; all ROIs contained a mixture of both +ve and −ve pRFs with neither an obvious spatial topography within an ROI, nor overall differences in explained variance (paired *t*-test of mean R^2^ *p*>0.05, in all cases except lhPlaces1, *p*=0.03). Next, we sought to test whether the pattern of visuospatial coding between +ve/−ve pRFs within each region could structure the responses during memory recall.

### Similarity in visuospatial coding between +ve/−ve pRFs structures responses during memory recall in MPC

Recent work identified a push-pull relationship between scene-perception and scene-memory areas of the brain during memory recall, such that when activity in +ve pRFs within scene-perception areas was high, activity in −ve pRFs within scene-memory areas was low and vice-versa - a relationship that was strengthened by the similarity in visuospatial coding between +ve/−ve pRF pairs (Steel et al., 2024). Crucially, this analysis was conducted ***between-regions***. This followed logically from the anatomical yoking of scene-perception and scene-memory areas on the lateral and ventral surfaces (Steel et al., 2021, 2024). An important difference between this perception-memory yoking and MPC is that within MPC there is not a clear gradient from perception-memory for both categories. Also, as demonstrated above, the prevalence of −ve pRFs did not change significantly throughout MPC (i.e., no effect of Position). Given this, we took a different ***within-region*** approach and asked whether the similarity in visuospatial coding between +ve/−ve pRFs within a region structures the responses during memory recall.

We start by considering a theoretical relationship between two pRFs within a region that are perfectly overlapping, but with opposite amplitudes (i.e., one +ve and one −ve, **Figure 3A**). Here, the two pRFs have identical centre locations and sizes (**left panel, Figure 3A**) but differ in that one responds positively to a stimulus within its receptive field (+ve pRF) and the other responds negatively (−ve pRF) with a perfect negative correlation (**middle & right panels, Figure 3A**). This scenario reflects an idealised scenario, but we can look for a similar pattern in real data (accepting that real data will be noisier).

**Figure 3:**
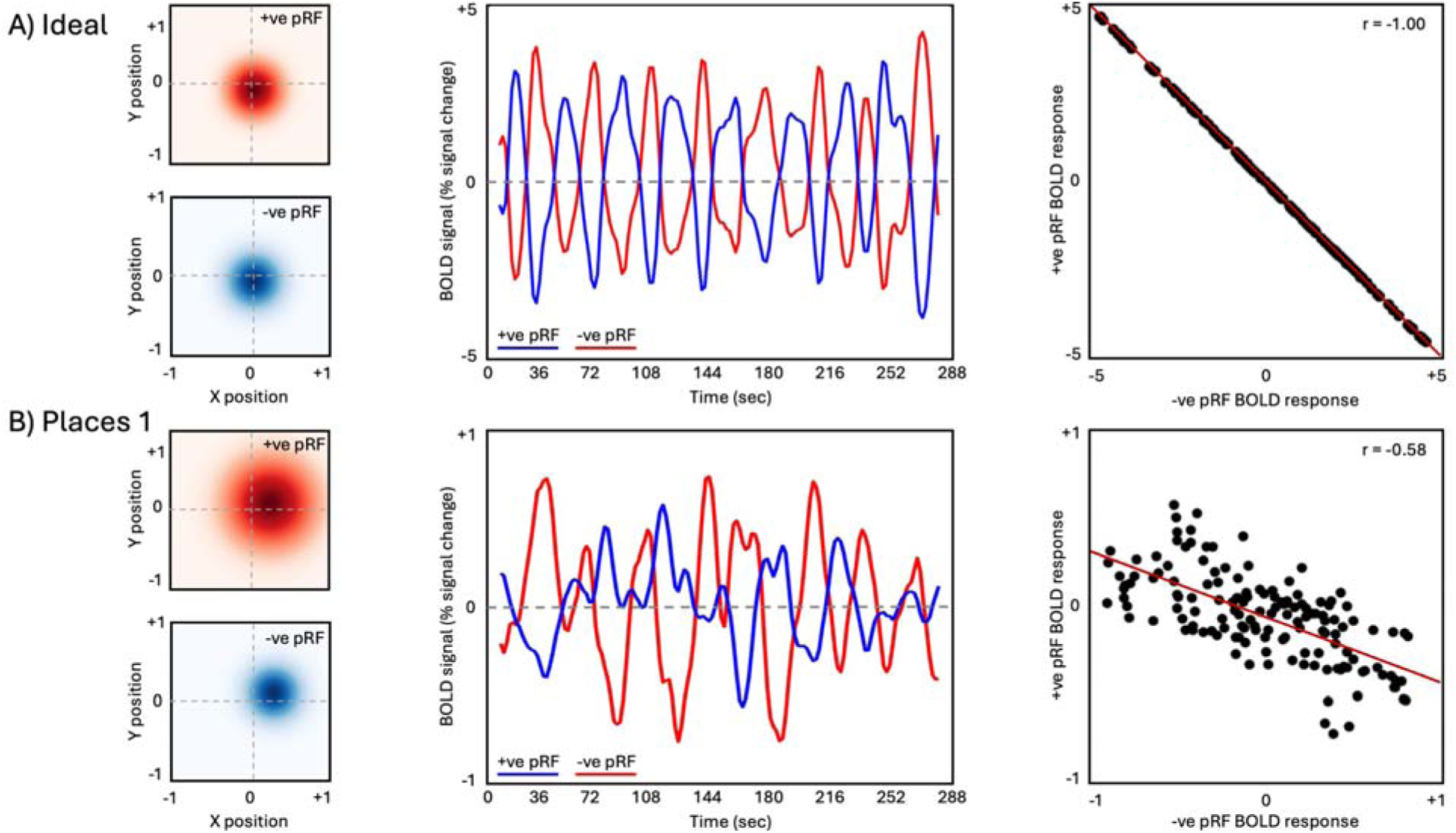
Opponent responses of matched +ve and −ve pRFs. **A) Left:** Idealised scenario of two perfectly overlapping pRFs with opposite amplitudes. **Middle:** The timeseries of the +ve pRF (red) and −ve pRF (blue) during pRF mapping. As activity in the +ve pRF increases in response to the bar stimulus within its receptive field, the activity of the −ve pRF decreases and vice-versa. **Right:** The two timeseries have a perfect negative correlation (r=-1.00). **B)** A pair of +ve/−ve pRFs from left hemisphere Places1 of a representative participant. These pRFs occup very similar portions of the visual field, although they are not perfectly overlapping. **Middle**: The timeseries of the +ve pRF (red) and −ve pRF (blue) during pRF mapping. As activity in the +ve pRF increases in response to the bar stimulus within its receptive field, the activity of the −ve pRF generally decreases. Note these timeseries have been smoothed for visualisation purposes. **Right:** The two timeseries are significantly negatively correlated (non-smoothed: r=-0.58, *p*<0.0001, smoothed: r=-0.66, *p*<0.0001).

**Figure 3B** depicts an example of two pRFs (one +ve and one −ve) from left hemisphere Places1 of a representative participant. Here, the +ve and −ve pRFs occupy very similar positions in the visual field (albeit not perfectly overlapping, **left panel Figure 3B**). Visualising the timeseries of these pRFs highlights the different patterns of response to the same visual stimulus within the +ve pRF (red-line) and −ve pRF (blue-line), respectively (**middle panel, Figure 3B**). There is a significant negative correlation between the responses of these two pRFs (r(142)= −0.58, *p*<0.0001, **right panel, Figure 3B**).

This indicative example demonstrates the inverse relationship expected between the timeseries during pRF mapping and the visual field coverage of +ve/−ve pRF pairs. That is, the more dissimilar the timeseries during pRF mapping between a pair of +ve and −ve pRFs, the more similar their coverage. As a proof of principle, we computed representational dissimilarity matrices (RDMs) that reflect the pairwise dissimilarity (1-Pearson’s r) between **1)** the pRF timeseries and **2)** the visual field coverage for all pairs of +ve and −ve pRFs within each participant and ROI. As anticipated, there exists on average a significant negative correlation between these two RDMs across participants for all ROIs (all *t*-values > 2.66, all *p*-values <0.02, **see Table S2 for full statistical breakdown**). This result demonstrates that the RDM of the pRF timeseries captures well the pattern of visuospatial coding between +ve and −ve pRFs within each MPC region.

Having established this relationship, we next assessed whether that same pattern of visuospatial coding structures the responses during memory recall. Here, we computed RDMs for each run of the recall task separately before averaging across runs in each participant and ROI (**see Methods**). These RDMs reflect the pairwise dissimilarity in response during recall between all pairs of +ve and −ve pRFs. Crucially then, if the dissimilarity in visuospatial coding between +ve and −ve pRFs (i.e., pRF timeseries RDM) within a region structures the responses during memory recall, then we expected positive correlations between these two RDMs across participants. **Figure 4** depicts this relationship for left hemisphere Places1 of a representative participant. The pattern of dissimilarity in visuospatial coding between +ve/−ve pRFs (**Figure 4A**) and the pattern of dissimilarity in memory recall response (**Figure 4B**) are positively correlated (**Figure 4C**). These data highlight that the more dissimilar the timeseries between a pair of +ve/−ve pRFs during pRF mapping, the more dissimilar their timeseries during memory recall - a task that places very different demands on this region.

**Figure 4:**
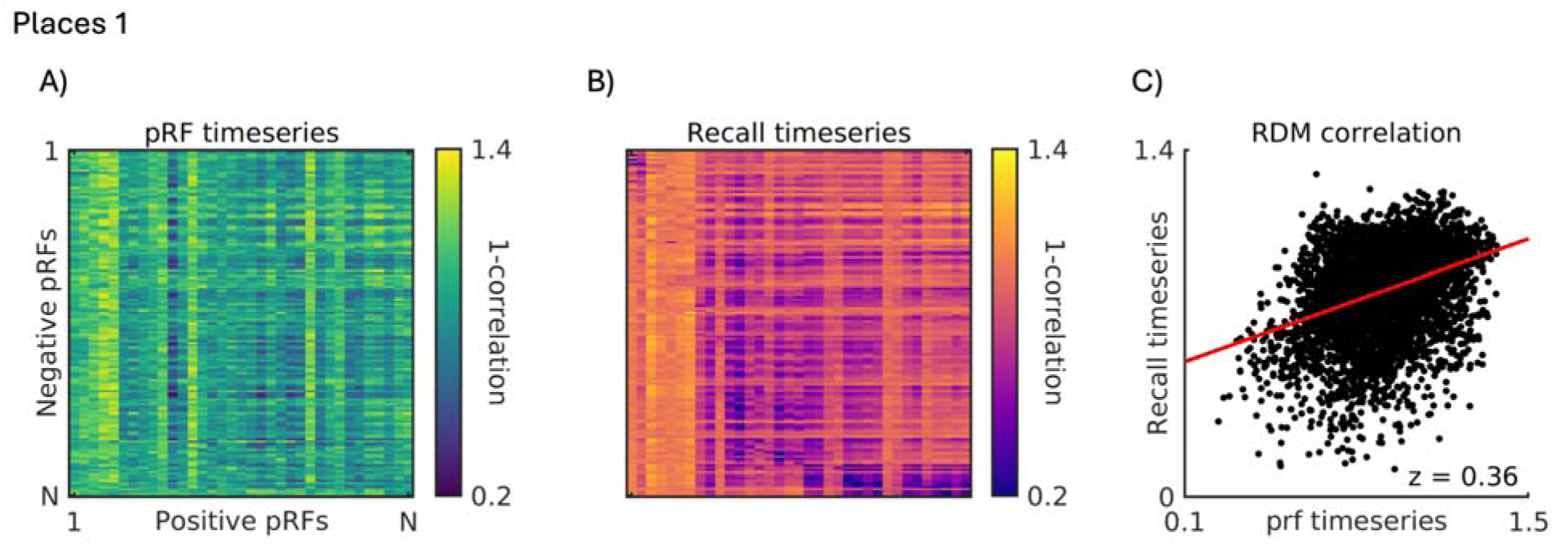
Positive relationship between pattern of visuospatial coding and memory recall response: **A)** An RDM representing the pattern of dissimilarity (1-Pearson’s r) in the timeseries of all +ve and −ve pRFs during pRF mapping (navy=more similar, yellow=more dissimilar). **B)** An RDM representing the average pattern of dissimilarity (1-Pearson’s r) in the timeseries of all +ve and −ve pRFs during memory recall (purple=more similar, yellow=more dissimilar). **C)** A scatter plot depicting the positive relationship between these two RDMs (z = 0.36).

As predicted, there exists on average significant and positive correlations between the pattern of dissimilarity in the pRF timeseries and the pattern of dissimilarity in memory recall between+ve/−ve pRFs within each ROI (all *t*-values > 2.87, all *p*-values <0.015, **Figure 5 & Table S3 fo full statistical breakdown**). This finding indicates that +ve and −ve pRFs with similar visuospatial coding (i.e., dissimilar timeseries but similar visual field coverage) within an ROI may represent different patterns of information during memory recall.

**Figure 5:**
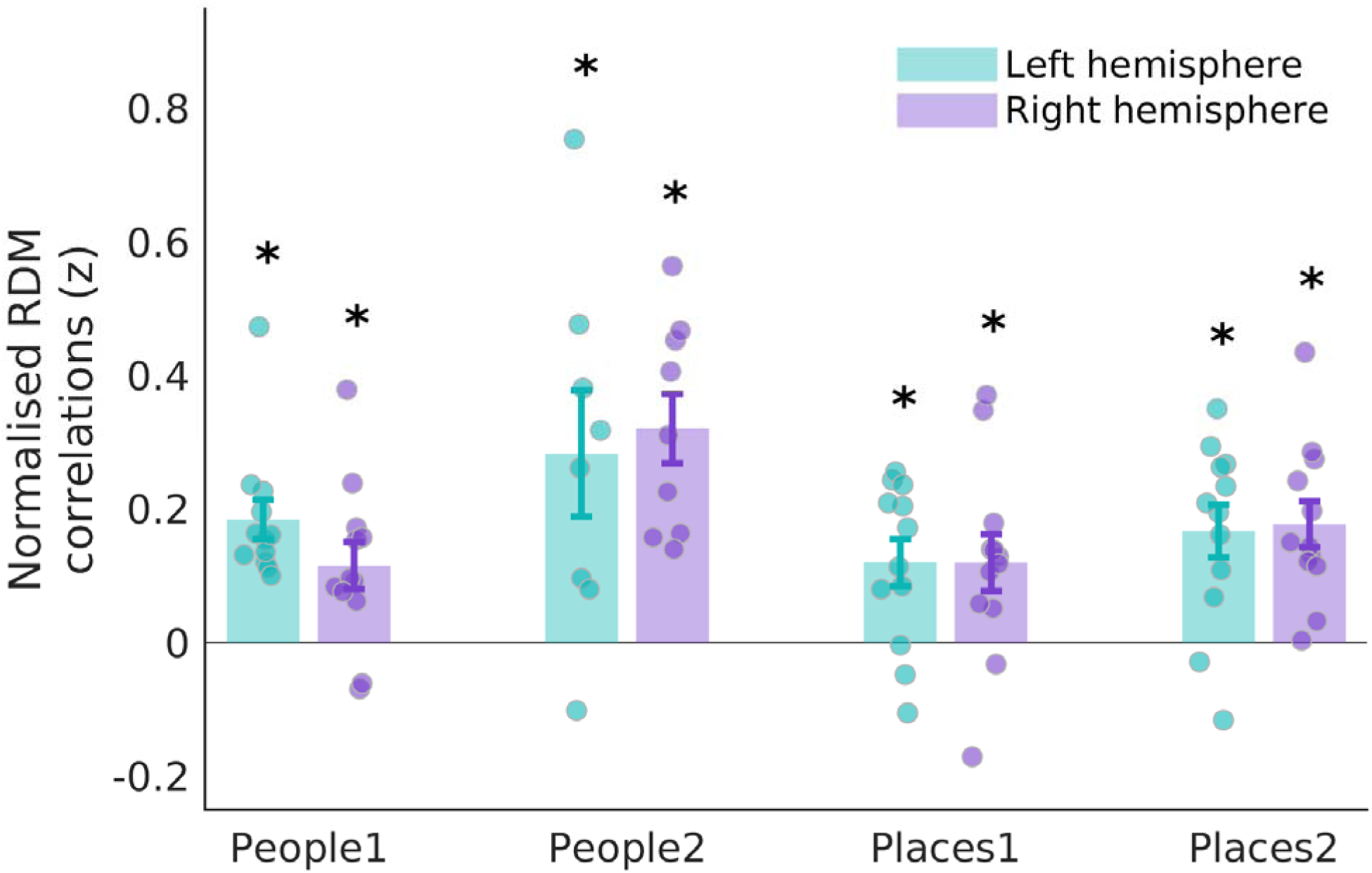
Pattern of visuospatial coding between +ve/−ve pRFs structures responses during memory recall. Bars represent the mean normalised correlation coefficients between the pRF timeseries RDM and memory recall RDM in each MPC ROIs (left hemisphere=green, right hemisphere=purple). Each data point represented an individual participant. Error bars represent SEM. The pattern of visuospatial coding between +ve/−ve pRFs was positively correlated with the pattern of responses during memory recall in each ROI (\**p*<0.05).

Is this relationship really structured by the similarity in visuospatial coding, or could it be due to less interesting differences like overall signal? To test this possibility, we performed two control analyses (**see Tables S4 & S5 for full statistical breakdown**). First, we computed the pairwise dissimilarity in temporal signal-to-noise-ratio (tSNR) between all +ve and −ve pRFs for both pRF and recall data in each participant and ROI. If the positive relationship observed above between the pRF and recall RDMs reflect overall signal differences, rather than the similarity in visuospatial coding, then we anticipated these tSNR RDMs to be positively correlated. On average, we observed no significant correlations between the pRF and recall tSNR RDMs (all *t*-values < 1.61, all *p*-values > 0.13, **see Table S4**). Second, we tested whether the pattern of responses during recall were really structured by the pattern of visuospatial coding by shuffling the labels of the +ve and −ve pRFs prior to computing the RDMs (shuffled RDMs). Shuffling the pRF labels destroys the similarity in visuospatial coding between +ve/−ve pRFs, and so if the visuospatial coding does indeed structure the responses during recall, then this shuffling should not produce any significantly positive correlations. As predicted, removing the visuospatial relationship between +ve and −ve pRFs destroyed the relationship between the shuffled pRF and recall RDMs (all *t*-values < 0.5, all *p*-values > 0.44, expect for rh Places2, which was significantly negative, **see Table S5**). Taken together, these control analyses confirm that it is the pattern of similarity in visuospatial coding between +ve/−ve pRFs that structures the pattern of response during memory recall.

### Similarity in visuospatial coding between +ve/−ve pRFs structures responses during visual perception

Our main analyses presented above focused on memory recall, given the recruitment of MPCs during mental imagery and the higher prevalence of −pRFs in MPC over lateral and ventral memory regions. However, it is also possible that the similarity in visuospatial coding between +ve and −ve pRFs reflects a representational principle that extends beyond a single cognitive process. That is, despite our ROIs being principally engaged during memory recall, it is possible that the visuospatial coding between +ve and −ve pRFs might also structure responses during other cognitive tasks, such as visual perception.

To test this possibility, we used data from an second group of participants (n=24) who performed a slow event-related visual scene-perception task (**see Methods**). First, we computed RDMs for each ROI based on the pRF timeseries. Like above, these RDMs represented the pairwise dissimilarity in the pRF response between +ve and −ve pRFs. Next, we computed RDMs based on the event-related timeseries (i.e., visual perception). If the similarity in visuospatial coding between +ve and −ve pRFs reflects a broad representational principle that structures the responses across different cognitive tasks, then we expected to find positive correlations between the pRF RDMs and the perceptual RDMs. **Figure 6** depicts this analysis for Places1 of a representative participant. As above, the similarity in visuospatial coding between +ve/−ve pRFs (**Figure 6A**) and the responses during visual perception (**Figure 6B**) were positively correlated (**Figure 6C**).

**Figure 6:**
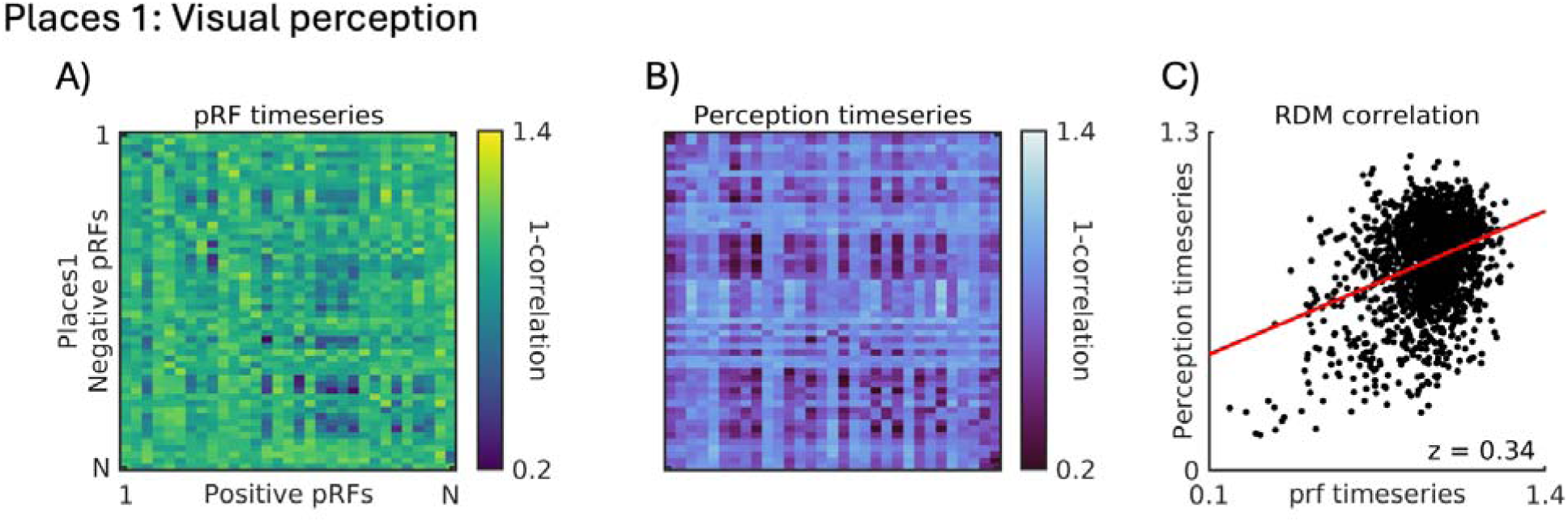
Positive relationship between pattern of visuospatial coding and pattern during visual perception. **A)** An RDM representing the pattern of dissimilarity (1-Pearson’s r) in the timeseries of all positive (+ve) and negative (−ve) pRFs during pRF mapping (navy=more similar, yellow=more dissimilar). **B)** An RDM representing the average pattern of dissimilarity (1-Pearson’s r) in the timeseries of all positive and negative pRFs during visual perception (dark blue=more similar, light blue=more dissimilar). **C)** A scatter plot depicting the positive relationship between these two RDMs (z = 0.34).

As anticipated, we observed significant and positive correlations between the pattern of dissimilarity in the pRF timeseries and the pattern of dissimilarity in the responses during visual perception (all *t*-values > 3.88, all *p*-values <0.001, **see Table S6**) between +ve/−ve pRFs within each ROI (**Figure 7**). This suggests that the visuospatial similarity between +ve and −ve pRFs is not simply a feature specific to pRF modelling, but extends to structure the relationship between voxels across different cognitive tasks.

**Figure 7:**
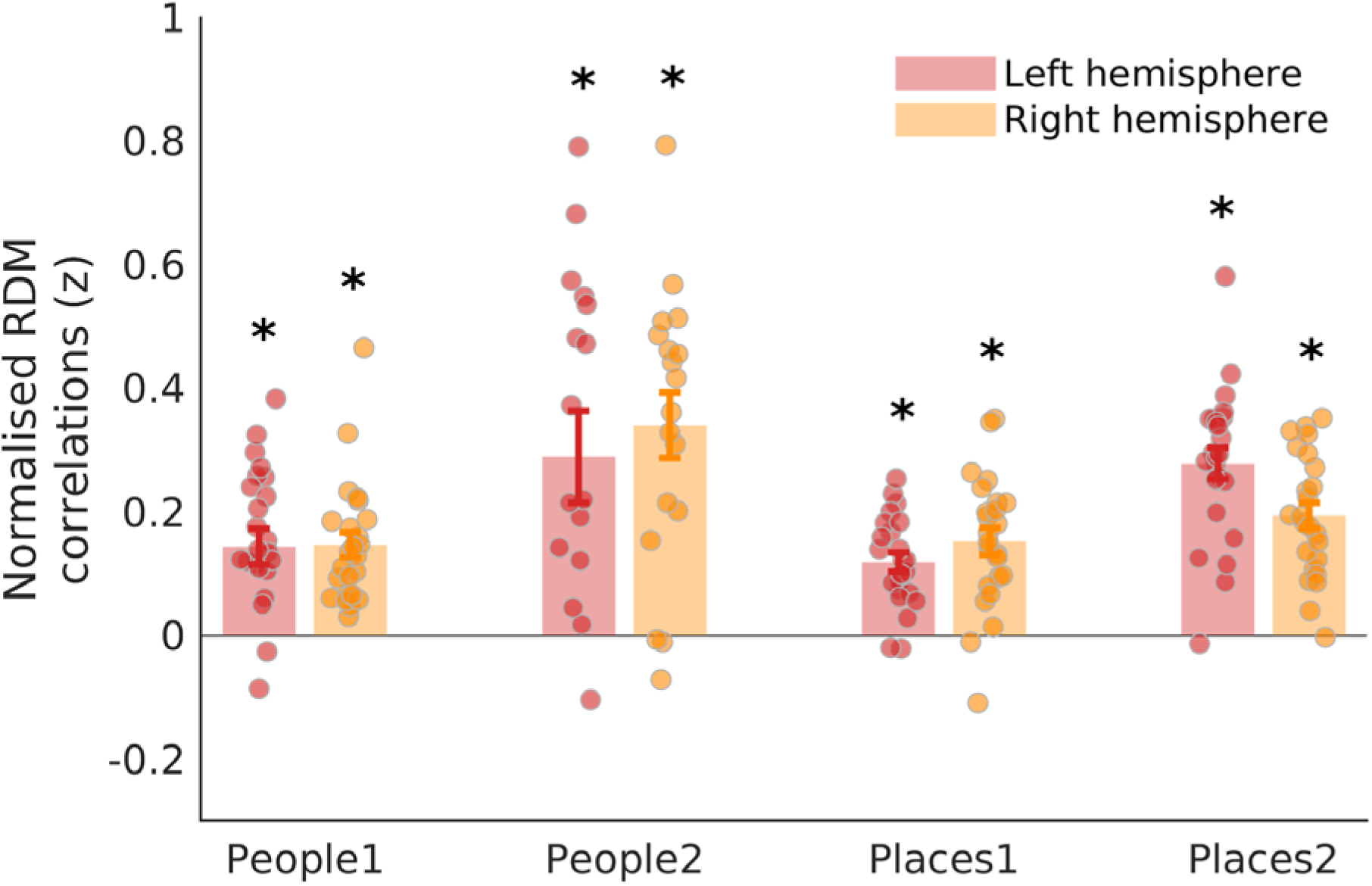
Pattern of visuospatial coding between +ve/−ve pRFs structures responses during visual perception recall. Bars represent the mean normalised correlation coefficients between the pRF timeseries RDM and visual perception RDM in each MPC ROIs (left hemisphere=red, right hemisphere=orange). Each data point represented an individual participant. Error bars represent the standard error of the mean. The pattern of visuospatial coding between +ve/−ve pRFs was positively correlated with the pattern of responses during visual perception in each ROI (\**p*<0.05).

## Discussion

We observed that the similarity in visuospatial coding between pRFs with opposite amplitudes structures their responses across different cognitive tasks, here within people and place memory areas of MPC. These data add to the growing body of literature demonstrating pervasive visuospatial coding throughout the brain (Groen et al., 2021), including visual cortex (Arcaro & Livingstone, 2017, 2021; Grill-Spector & Weiner, 2014; Silson et al., 2015; Wandell et al., 2007), the default mode network (Szinte & Knapen, 2020) and hippocampus (Knapen, 2021; Silson, Zeidman, et al., 2021). Our data extends recent work (Steel et al., 2024) by showing that visuospatial coding is present within people and place memory areas within MPC (Silson, Steel, et al., 2019).

Prior work using data from the Human Connectome Project initiative first identified −ve pRFs within the default-mode-network (Andrews-Hanna et al., 2010; Raichle et al., 2001), including its MPC component (Szinte & Knapen, 2020). The authors speculated on, but did not demonstrate, the potential functional significance of these −ve pRFs. Within the context of MPC, we replicate this finding (Szinte & Knapen, 2020) by showing significant levels of −ve pRFs within people and place recall regions. Moreover, we demonstrate that the relationship between +ve/−ve pRFs is not simply an inherited feature of the pRF modelling technique but persists during tasks that place very different demands on the default-mode-network (i.e., internally oriented memory recall versus externally oriented visual perception).

The functional role of MPC is often considered within the context of the default-mode-network (Andrews-Hanna et al., 2010; Andrews-Hanna et al., 2014; Raichle et al., 2001; Shirer et al., 2012), where much of MPC is thought to act a as a ‘core’ region that integrates information between separate dorsal and ventral subnetworks (Andrews-Hanna et al., 2010). The original delineation of people and place recall areas within MPC (Silson, Steel, et al., 2019) challenged this conceptualisation by suggesting that this core itself is fractionated along the same lines as the dorsal and ventral subnetworks; Places1/2, like the ventral component, are associated with scene-construction or contextual association, whereas People1/2, like the dorsal component, are associated with more social and potentially semantic processing (Andrews-Hanna et al., 2010; Bar & Aminoff, 2003; Hassabis et al., 2007). The current data add further complexity to this picture by highlighting that not only do these regions contain a mixture of +ve and −ve visually sensitive pRFs, but that the relationship between pairs of +ve/−ve pRFs within a region structures the responses of those voxels more generally.

Why do these regions contain a mixture of +ve/−ve pRFs? One possibility (already discussed in the context of −ve pRFs in the default-mode-network) is that there are computational benefits to representing the same signal with both activation and deactivation (Szinte & Knapen, 2020). For example, efficiency of sensory processing can be increased through the interplay of activation and deactivation via predictive coding (Srinivasan et al., 1982), and deactivations can help discount erroneous computational outcomes by explicitly representing ‘what is not’ (Goncalves & Welchman, 2017). The fact that each region contains a mixture of +ve/−ve pRFs offers the possibility that such an interplay occurs in MPC. For example, one interpretation of −ve pRFs was that they could serve to store perceptual signals before they are used for other forms of cognition (Groen et al., 2021; Szinte & Knapen, 2020). The current data are consistent with this idea, but suggest that such signals are potentially stored and accessed within a region. That responses during both memory recall and visual perception become more dissimilar the more similar a pair of +ve/−ve pRFs become in visuospatial terms hints strongly at just such an interplay.

The finding that the more similar a pair of +ve and −ve pRFs are in terms of visuosaptial coding the more dissimilar their responses during different cognitive tasks is consistent with the idea that the default-mode-network is well situated to integrate information across different functional modalities (Margulies et al., 2016; Popham et al., 2021b). Matched visual and language areas have been identified recently at the anterior border of visual cortex (Popham et al., 2021b). Within MPC specifically, two regions in the approximate locations of Places1 and People1 were found to have aligned visual and semantic representations. It is possible that such alignment across different representations is facilitated via the opponent dynamics of +ve/−ve pRFs within these regions, in the same way that responses during memory recall and visual perception were structured via these interactions reported here. Considering this prior work, it is interesting to note that although the regions we describe as Places1 and People1 appear commensurate with regions showing aligned visual and semantic representations (Popham et al., 2021b), these do not extend anteriorly to cover the regions we describe as Places2 and People2, suggesting perhaps that the more anterior pair of recall regions in MPC differ from their posterior counterparts on some functional level as yet not identified. Indeed, in our data we do not see clear evidence of a posterior-anterior shift in either the prevalence of −ve pRFs or in the degree to which the pattern of visuospatial coding relates to the recall or perceptual responses. Future work should aim to establish what (if any) functional component differentiates the posterior and anterior pairs of recall regions within MPC.

Several important distinctions between the current data and prior recent work (Steel et al., 2024) are noteworthy. First, the spatial scale of visuospatial coding interactions is very different. For instance, prior work observed a push-pull relationship between +ve and −ve pRFs in separate, albeit adjacent scene-perception and scene-memory areas, respectively (Steel et al., 2024). It was shown that as activity in +ve pRFs increased, so activity in −ve pRFs decreased and vice-versa. This visuospatial interaction occurred therefore at the ***between-region*** level. Here, we observe ***within-region*** interactions between +ve and −ve pRFs that structure the responses of those voxels across multiple cognitive tasks. Second, prior work focused exclusively on scene-perception and scene-memory areas on the lateral and ventral surfaces and on responses during scene-perception and scene-memory, respectively. Here, we extend the scope of visuospatial coding to include both people and place memory regions of MPC and measure responses during memory recall of people and places, as well as during visual perception.

Taken together, these data add to the growing body of work demonstrating the pervasive influence of visuospatial coding throughout the brain. Within MPC specifically, we show that not only do people and place memory regions exhibit robust signatures of visuospatial coding in the form of +ve and −ve pRFs, but that the similarity in visuospatial representation between +ve/−ve pRFs structures the similarity in response during both internally-orientated memory recall and externally-orientated visual perception. The interplay between +ve/−ve pRFs in MPC may play a vital role in integrating information across different representational spaces.

## Data and Code Availability

Data and code will be made available via the Open Science Framework upon acceptance.

## Author Contributions

CS & EHS jointly conceived of the project. CS collected the data. CS & ES jointly analyzed the data and wrote the manuscript.

## Supplementary Material

**Table S1:**
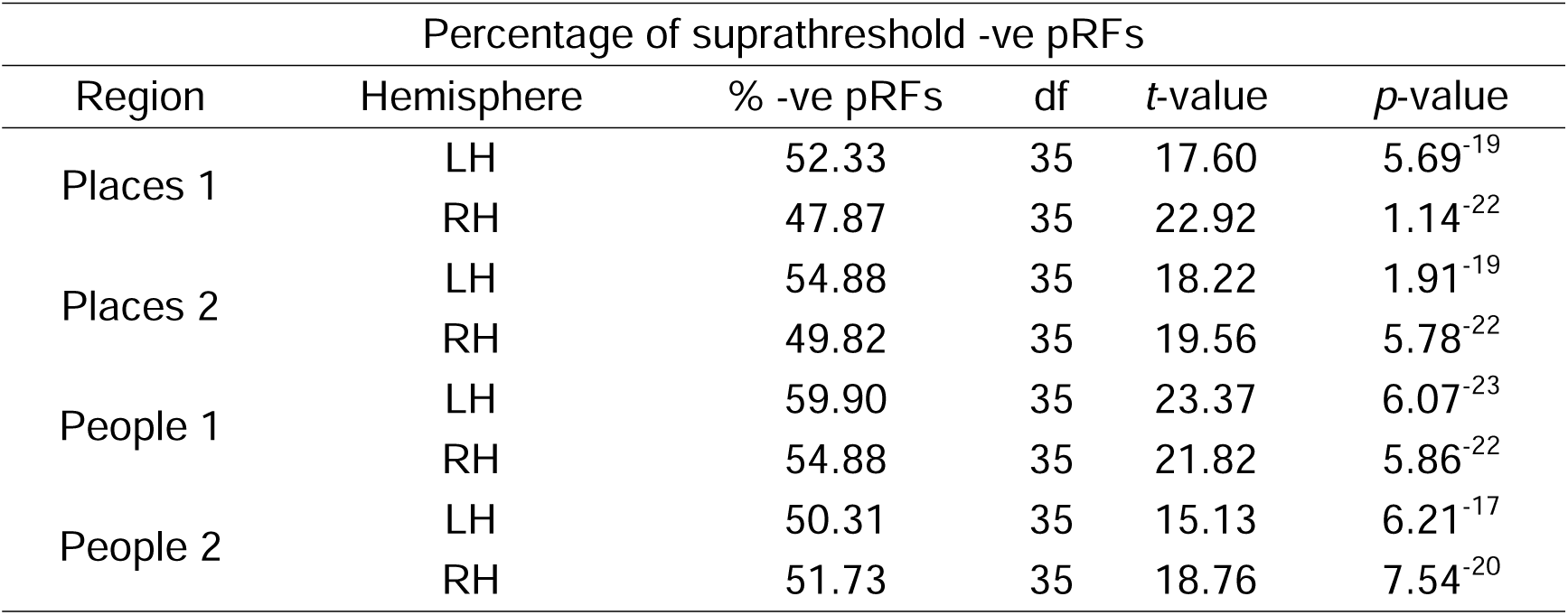
Table contains the percentage of supreahtrehsold −ve pRFs in each ROI, degrees of freedom (df), *t*-values and *p*-values for the two-tailed *t*-tests against zero.

**Table S2:**
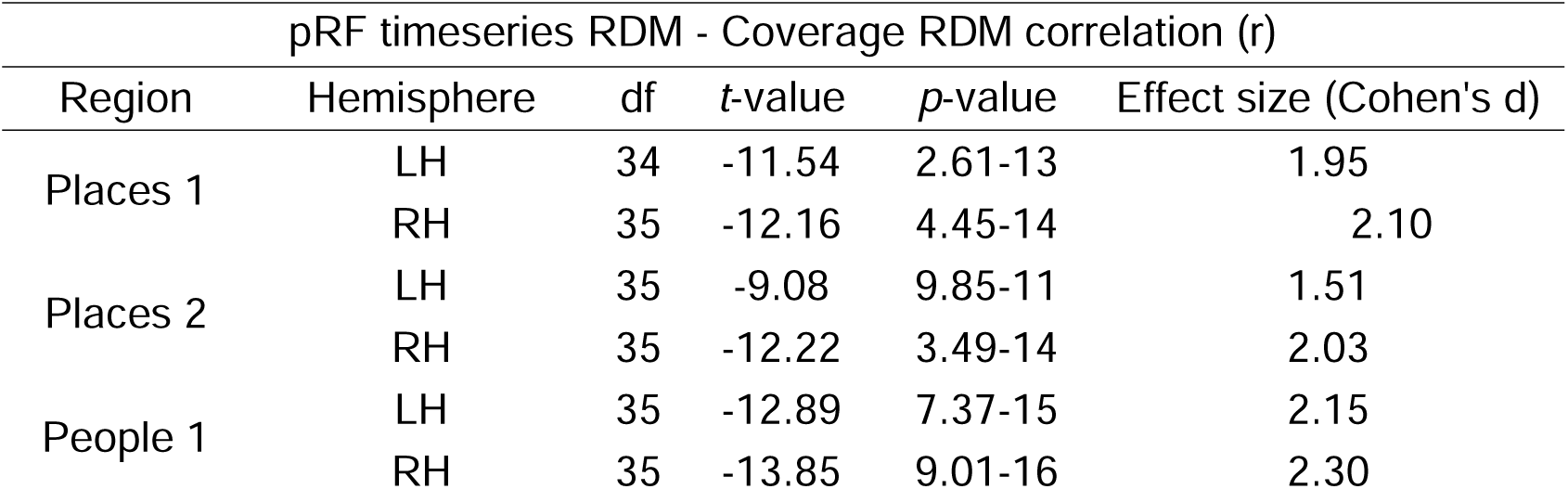
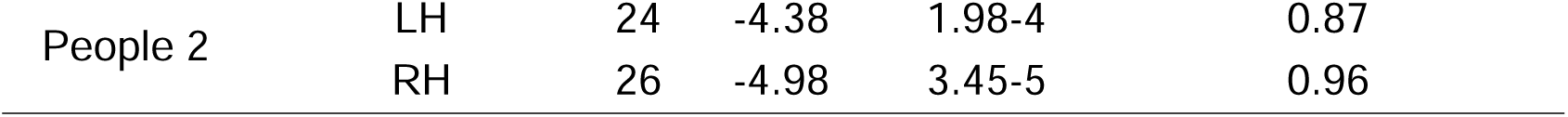
Table contains the degrees of freedom (df), *t*-values, *p*-values and effect size estimates for the correlation between pRF timeseries RDM and Coverage RDM for each ROI.

**Table S3:** Table contains the degrees of freedom (df), *t*-values, *p*-values and effect size estimates for the correlation between pRF timeseries RDM and Recall RDM for each ROI.

| pRF timeseries RDM - Recall RDM correlation (r) |  |  |  |  |  |
| --- | --- | --- | --- | --- | --- |
| Region | Hemisphere | df | *t*-value | *p*-value | Effect size (Cohen's d) |
| Places 1 | LH | 11 | 3.44 | 0.005 | 1.01 |
|  | RH | 11 | 2.84 | 0.016 | 0.82 |
| Places 2 | LH | 11 | 4.23 | 0.001 | 1.22 |
|  | RH | 11 | 5.19 | 2.97-4 | 1.49 |
| People 1 | LH | 11 | 6.35 | 5.40-5 | 1.83 |
|  | RH | 11 | 3.29 | 0.007 | 0.95 |
| People 2 | LH | 7 | 3.01 | 0.01 | 1.06 |
|  | RH | 8 | 6.11 | 2.84-4 | 2.03 |

**Table S4:**
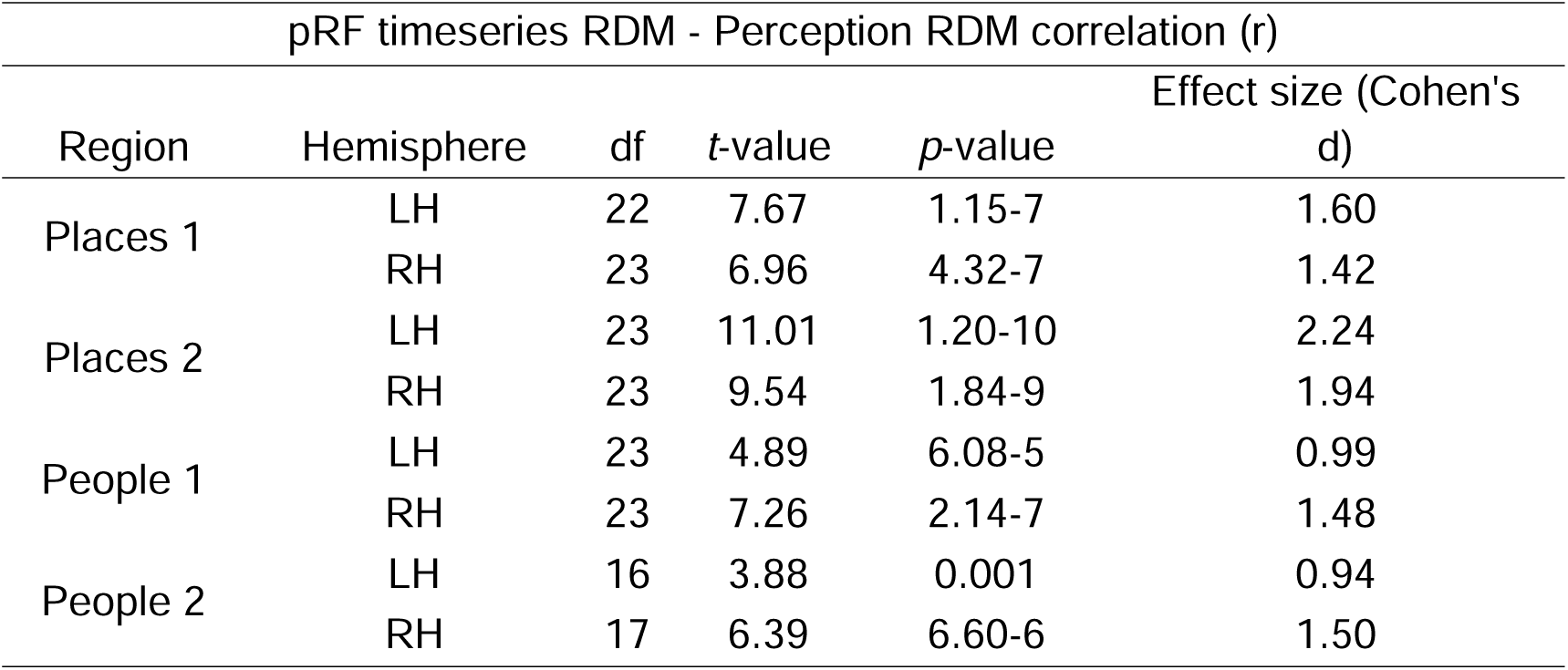
Table contains the degrees of freedom (df), *t*-values, *p*-values and effect size estimates for the correlation between pRF timeseries RDM and Perception RDM for each ROI.

**Table S5:** Table contains the degrees of freedom (df), *t*-values, *p*-values and effect size estimates for the correlation between pRF tSNR and Recall tSNR RDMs for each ROI.

| pRF tSNR RDM – Recall tSNR RDM correlation (r) |  |  |  |  |  |
| --- | --- | --- | --- | --- | --- |
| Region | Hemisphere | df | *t*-value | *p*-value | Effect size (Cohen's d) |
| Places 1 | LH | 11 | 1.61 | 0.31 | 0.30 |
|  | RH | 11 | 0.16 | 0.87 | 0.04 |
| Places 2 | LH | 11 | 0.21 | 0.83 | 0.06 |
|  | RH | 11 | 0.79 | 0.44 | 0.22 |
| People 1 | LH | 11 | 1.06 | 0.31 | 0.30 |
|  | RH | 11 | 0.46 | 0.65 | 0.13 |
| People 2 | LH | 7 | 0.45 | 0.66 | 0.16 |
|  | RH | 8 | 0.29 | 0.77 | 0.09 |

**Table S6:** Table contains the degrees of freedom (df), *t*-values, *p*-values and effect size estimates for the correlation between the shuffled pRF timeseries and Recall RDMs for each ROI.

| Shuffled pRF RDM – shuffled Recall RDM correlation (r) |  |  |  |  |  |
| --- | --- | --- | --- | --- | --- |
| Region | Hemisphere | df | *t*-value | *p*-value | Effect size (Cohen's d) |
| Places 1 | LH | 11 | 0.28 | 0.78 | 0.08 |
|  | RH | 11 | -2.72 | 0.02 | 0.78 |
| Places 2 | LH | 11 | -0.80 | 0.44 | 0.23 |
|  | RH | 11 | 0.55 | 0.59 | 0.16 |
| People 1 | LH | 11 | 0.11 | 0.91 | 0.03 |
|  | RH | 11 | 0.62 | 0.54 | 0.17 |
| People 2 | LH | 7 | -0.49 | 0.63 | 0.17 |
|  | RH | 8 | -0.36 | 0.72 | 0.16 |

